# Molecular diversity of glutamatergic and GABAergic synapses from multiplexed fluorescence imaging

**DOI:** 10.1101/2020.06.12.148155

**Authors:** Eric Danielson, Karen Perez de Arce, Beth Cimini, Eike-Christian Wamhoff, Shantanu Singh, Jeffrey R Cottrell, Anne E. Carpenter, Mark Bathe

**Affiliations:** Department of Biological Engineering, MIT, Cambridge, MA, USA; Stanley Center for Psychiatric Research, Broad Institute of MIT and Harvard, Cambridge, MA, USA; Imaging Platform, Broad Institute of MIT and Harvard, Cambridge, MA, USA

## Abstract

Neuronal synapses contain hundreds of different protein species important for regulating signal transmission. Characterizing differential expression profiles of proteins within synapses in distinct regions of the brain has revealed a high degree of synaptic diversity defined by unique molecular organization. Multiplexed imaging of *in vitro* neuronal culture models at single synapse resolution offers new opportunities for exploring synaptic reorganization in response to chemical and genetic perturbations. Here, we combine 12-color multiplexed fluorescence imaging with quantitative image analysis and machine learning to identify novel synaptic subtypes within excitatory and inhibitory synapses based on the expression profiles of major synaptic components. We characterize differences in the correlated expression of proteins within these subtypes and we examine how the distribution of these synapses is modified following induction of synaptic plasticity. Under chronic suppression of neuronal activity, phenotypic characterization revealed coordinated increases in both excitatory and inhibitory protein levels without changes in the distribution of synaptic subtypes, suggesting concerted events targeting glutamatergic and GABAergic synapses. Our results offer molecular insight into the mechanisms of synaptic plasticity.

## Introduction

Synapses contain complex proteomes that organize into multiprotein signaling complexes (Collins et al. 2006; Husi et al. 2000). There appears to be a high degree of diversity in the expression, stoichiometry, and organization of these proteins across scales from individual synapses to the entire brain (Roy et al. 2018; Zhu et al. 2018; Richard D. Emes and Grant 2012). While it is unknown how many classes of synapses exist, there are over 1,000 genes that encode synaptic proteins (Husi et al. 2000; Peng et al. 2004; Collins et al. 2005; 2006; Richard D Emes et al. 2008; Bayés et al. 2011; 2012; 2017; Richard David Emes and Grant 2011; Distler et al. 2014). The differential expression of these proteins across distinct brain regions, as well as differential spatial-temporal expression during development, suggest significant synaptic diversity. Immunofluorescence labeling of two differentially expressed excitatory scaffolding proteins was used to examine synaptic diversity throughout the mouse brain (Zhu et al. 2018) however, to date how synaptic diversity is affected by synaptic scaling has not been investigated. Synaptic scaling maintains neuronal homeostasis via global changes in expression of multiple synaptic proteins targeting both excitatory and inhibitory synapses to prevent runaway neuronal excitability (G. G. Turrigiano 2008; G. Turrigiano 2012; G. G. Turrigiano and Nelson 2000; 2004; G. G. Turrigiano et al. 1998). Examination of the entire molecular composition of individual synapses using techniques that can survey the entire proteome, such as mass spectrometry, would be ideal for characterizing synapses. However, synapses are approximately 2 μm in size; and while advances in mass spectrometry-based imaging have achieved 1 μm resolution (Zavalin et al. 2015), the majority of commercial matrix-assisted laser desorption/ionization (MALDI) mass spectrometers are not sufficiently accurate for examination of individual synapses. Hence, microscopy techniques, such as immunofluorescence, remains the optimal technique for the examination of individual synapses. However, conventional fluorescence microscopy is generally limited to four channels as a result of the maximal spectral resolution of organic and biomolecular fluorophores. This limitation presents a challenge to comprehensive analysis of synaptic architecture due to the large number of different protein species within each synapse.

Multiplexed imaging techniques are particularly useful for studying neuronal synapses, and investigating the coordinated assembly of dozens to hundreds of distinct proteins involved in synaptic development, function, and plasticity. Probe-based Imaging for Sequential Multiplexing (PRISM) is a recently introduced multiplexed imaging technique that uses single-stranded deoxyribonucleic acid (ssDNA)-conjugated antibodies or peptides with complementary fluorescently labeled single-stranded locked nucleic acid (ssLNA) imaging probes to sequentially visualize multiple synaptic targets *in situ* (Guo et al. 2019). With this approach, the affinity of the ssLNA imaging probes is salt-dependent, allowing multiple synaptic targets to be imaged within the same sample through sequential rounds of imaging first in physiological salt buffer, followed by rapid imaging-strand removal in low-salt buffer. This method of imaging prevents the neuronal sample disruption that occurs with alternative multiplex imaging techniques (Gerdes et al. 2013; Lin, Fallahi-Sichani, and Sorger 2015). PRISM has been used to quantify changes in synaptic protein levels, co-expression profiles, and synapse-subtype compositions following blockade of action potentials with tetrodotoxin (TTX) treatment (Guo et al. 2019). This analysis revealed that TTX induces a coordinated reorganization of excitatory presynaptic and postsynaptic proteins. However, structural plasticity in response to chronic activity changes occurs in both excitatory and inhibitory synapses, warranting an expansion of our previous analysis by incorporating inhibitory synapse imaging and analysis.

Here, we used PRISM to measure homeostatic structural changes of subcellular compartments of rat hippocampal neurons, simultaneously discriminating between excitatory and inhibitory terminals. We include the inhibitory markers vGAT and gephyrin in addition to a subset of excitatory markers previously analyzed by PRISM imaging (Guo et al. 2019). In addition, we created an automated imaging and analysis pipeline in the open-source bioimage analysis software CellProfiler that identifies and characterizes synaptic puncta from multiplexed images to compare changes in protein levels that occur in excitatory and inhibitory synapses. Using this computational framework, we identified six distinct classes of synapses and found differential regulation of synapsin1 at both excitatory and inhibitory synapses in response to TTX treatment. However, while we found increased inhibitory presynaptic protein levels following activity suppression, TTX treatment did not affect synaptic diversity in 21 days *in vitro* (DIV 21) neurons. These results broaden our knowledge of the molecular events that occur during synaptic plasticity.

## Results

### Characterization of synaptic content from multiplexed images

We designed a Cell Profiler pipeline to automate synapse detection and protein analysis. Analysis of individual synapses from multiplexed images requires three main computational steps: image alignment, synapse segmentation, and synapse alignment. To address these technical requirements, we generated a CellProfiler pipeline that imports multiplexed images (four single-channel images from a single field), aligns images from ten different imaging rounds of four channels using MAP2 staining, and uses the synapsin1 channel for synapse identification. Puncta signals are enhanced using a white top-hat filter and then segmented using intensity peaks (**Figure 1A and Supplemental Figure 1 and 2**). Following segmentation, each individual punctum from each channel is then assigned to a synapsin1 cluster, depending on the percentage of overlap with the synapsin1 signal (**see Methods**). Thus our “synapses” are a conglomeration of puncta: a single synapsin1 punctum and the collection of individual puncta from the other channels that overlap with that synapsin1 cluster. A large number of geometric and intensity feature calculations are then performed to enable a comprehensive and detailed phenotypic analysis (**Supplementary Table 1**). Specifically, we measured multiple intensity and shape features for each punctum to later identify synaptic clusters.

**Figure 1.**
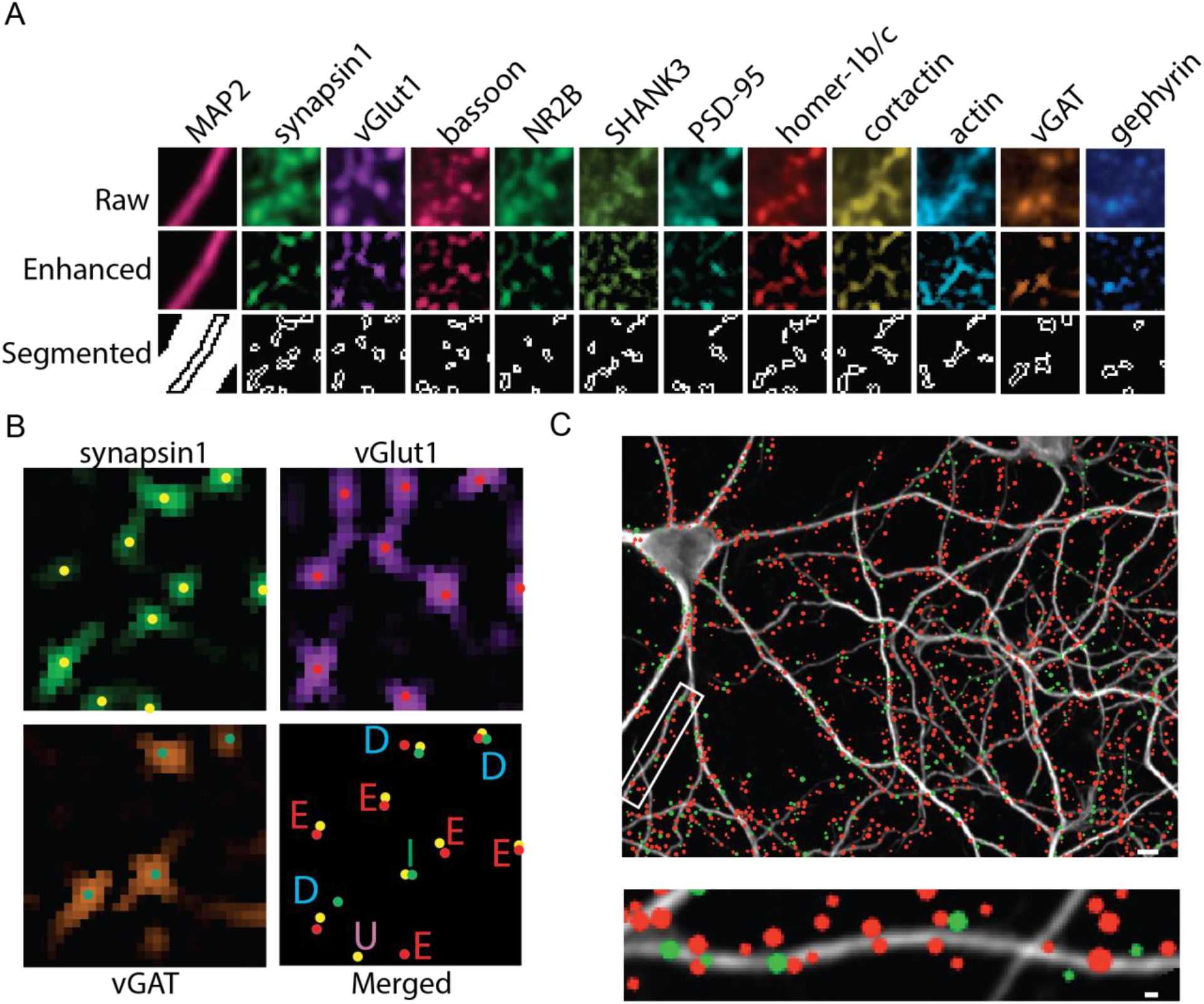
Measuring synaptic content from excitatory and inhibitory synapses using CellProfiler. (A) Multiplexed images of PRISM stained DIV 21 hippocampal neurons are used as input for the CellProfiler pipeline. Raw images (top) are aligned using MAP2 signal. A white-top filter is applied to enhance puncta (middle). Puncta are thresholded and separated into individual puncta using peak intensity (bottom). (B) Synapses are labeled excitatory (E) or inhibitory (I) using the presence or absence of excitatory specific (vGlut1) or inhibitory specific (vGAT) markers. Synapses with both markers are labeled as dual (D). Clusters that do not contain either marker are labeled as unknown (U). (C) Representative image (top) and enlarged dendrite (bottom) from a MAP2 stained DIV 21 hippocampal neuron with excitatory (red) and inhibitory (green) synapses labeled as colored circles. The size of the colored circle represents the relative synapsin1 area. Scale bar 1 μm.

Primary rat hippocampal neurons were grown for 19 days in culture, then fixed and stained with our PRISM antibodies/peptides: PSD-95, synapsin1, bassoon, actin, cortactin, homer-1b/c, SHANK3, NR2B, and vGlut1 plus two antibodies against inhibitory synaptic proteins, vGAT and gephyrin (**Figure 1A**). Synapses were defined as excitatory when the presynaptic proteins synapsin1 and vGlut1 were present versus inhibitory when synapsin1 and vGAT were present (**Figure 1C and D**). Under this classification criterion, 66% of detected puncta were defined as synapses and used for further characterization (**Supplemental Figure 3**). Synapses that failed to meet our criteria of classification for glutamatergic or GABAergic synapses were excluded from the analysis. These types of puncta were labeled as unclassified (**Supplemental Figure 3**) because they either lacked vGlut1 or vGAT (~22%, ‘Unknown’), or contained both synaptic markers (~12%, ‘Dual’). Thus, the CellProfiler pipeline enabled us to restrict our analysis of synaptic composition to synapses we could confidently identify.

### Characterization of synapse diversity

To characterize synaptic diversity in hippocampal cultured neurons, Uniform Manifold Approximation and Projection (UMAP) was applied to a subset of the CellProfiler pipeline output (**Figure 2A**). In total, 15 separate intensity measurements, 14 separate punctum shape measurements, quantification of punctum distances to synapsin1, and quantification of the number of puncta associated with each synapse were measured using CellProfiler (**Supplementary Table 1**). Feature reduction (**see Methods**) was used to isolate the most important features prior to clustering. The UMAP output shows clearly distinct excitatory and inhibitory synapses, which were found at a ratio of 5 excitatory to 1 inhibitory, consistent with the ratio observed using the CellProfiler pipeline to classify synapses (**Figure 2A**). Most (98.9%) of the synapses present in the excitatory clusters were positive for vGlut1 staining whereas only 1.1% of the synapses in those clusters contained vGAT. Similarly, 97.3% of synapses present in the inhibitory cluster contained synapsin1 and vGAT and only 2.7% of these synapses contained synapsin1 and vGlut1.

**Figure 2.**
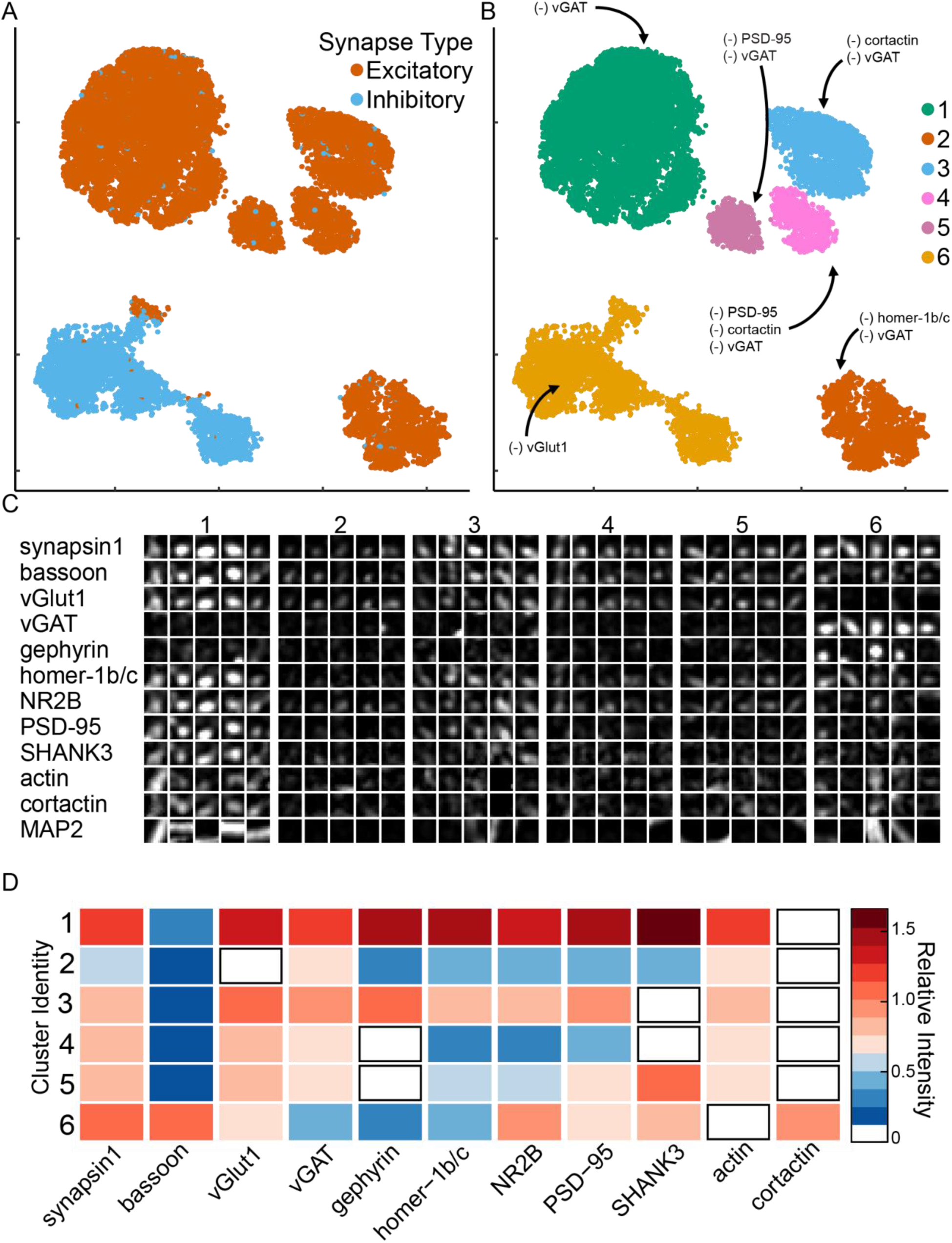
Synaptic cluster identification using UMAP. (A) UMAP plots of individual synapses (n = 10,000) using CellProfiler output separates excitatory (red) and inhibitory (blue) synapses into two major clusters with multiple sub-clusters. (B) Unique clusters identified by HDBSCAN. (-) indicates synaptic target below limit of detection. (C) Representative synapses from each cluster. (D) Heatmap indicates the average relative intensities for each synaptic target within each cluster. All values are normalized to untreated mean integrated intensity.

To evaluate the quality of the segmentation, we assessed whether canonical proteins specific to one type of synapse were present at excitatory and inhibitory synapses. We found the majority (78.8%) of the excitatory synapses did not contain the inhibitory target gephyrin and those that did express gephyrin had lower synaptic levels compared to inhibitory synapses (power analysis, effect = 0.51). Moreover, synapsin1 and gephyrin were further apart (mean distance 1.22+0.02 pixels) than synapsin1 and the excitatory marker PSD-95 (0.92+0.01 pixels) suggesting the detected gephyrin was likely outside of these excitatory synapses, but overlapping due to limited spatial resolution. Similarly, the excitatory marker PSD-95 was detectable in 35.7% of inhibitory synapses; however, synaptic levels of PSD-95 within the inhibitory cluster were reduced(power analysis, effect = 0.49) and the mean distance between synapsin1 and PSD-95 (1.5+0.04 pixels) was almost twice that of synapsin1 and gephyrin (0.8+0.02 pixels). Thus, given the limits of the imaging resolution and density of the neuronal culture, the image segmentation may have incorrectly assigned proteins to some synapses; however, the assignment of proteins to synapses can be further refined, as needed, using features such as protein levels and distance to synapsin1.

We examined the UMAP output in further detail to identify and characterize synapses with distinct protein expression profiles. Using Hierarchical Density-Based Spatial Clustering of Applications with Noise (HDBSCAN) applied to the UMAP output, we identified six unique clusters of synaptic subtypes (**Figure 2B, C)**. Subsequently, we generated heatmaps (**Figure 2D**) and additional scatterplots (**Supplemental Figure 4-16)** to distinguish differences among these clusters. Based on the relative vGlut1 and vGAT levels, excitatory synapses correspond to Clusters 1 to 5 and inhibitory synapses are contained in Cluster 6. Cluster 1 was the largest cluster with 57.9% of the excitatory synapses and had the highest levels of synaptic proteins for all targets relative to the other excitatory clusters (**Figure 2D and Supplemental Figure 17**). Compared to Cluster 1, Clusters 2-5 contain lower levels of all synaptic proteins and each of these clusters had very low levels of one or more postsynaptic scaffolding or cytoskeletal protein. To investigate in more detail the differences among the excitatory synaptic clusters and identify the distinguishing features for each synapse subtype, we measured the protein intensity profiles, correlation coefficients, and performed hierarchical clustering for each of the individual clusters (**Figure 3**).

**Figure 3.**
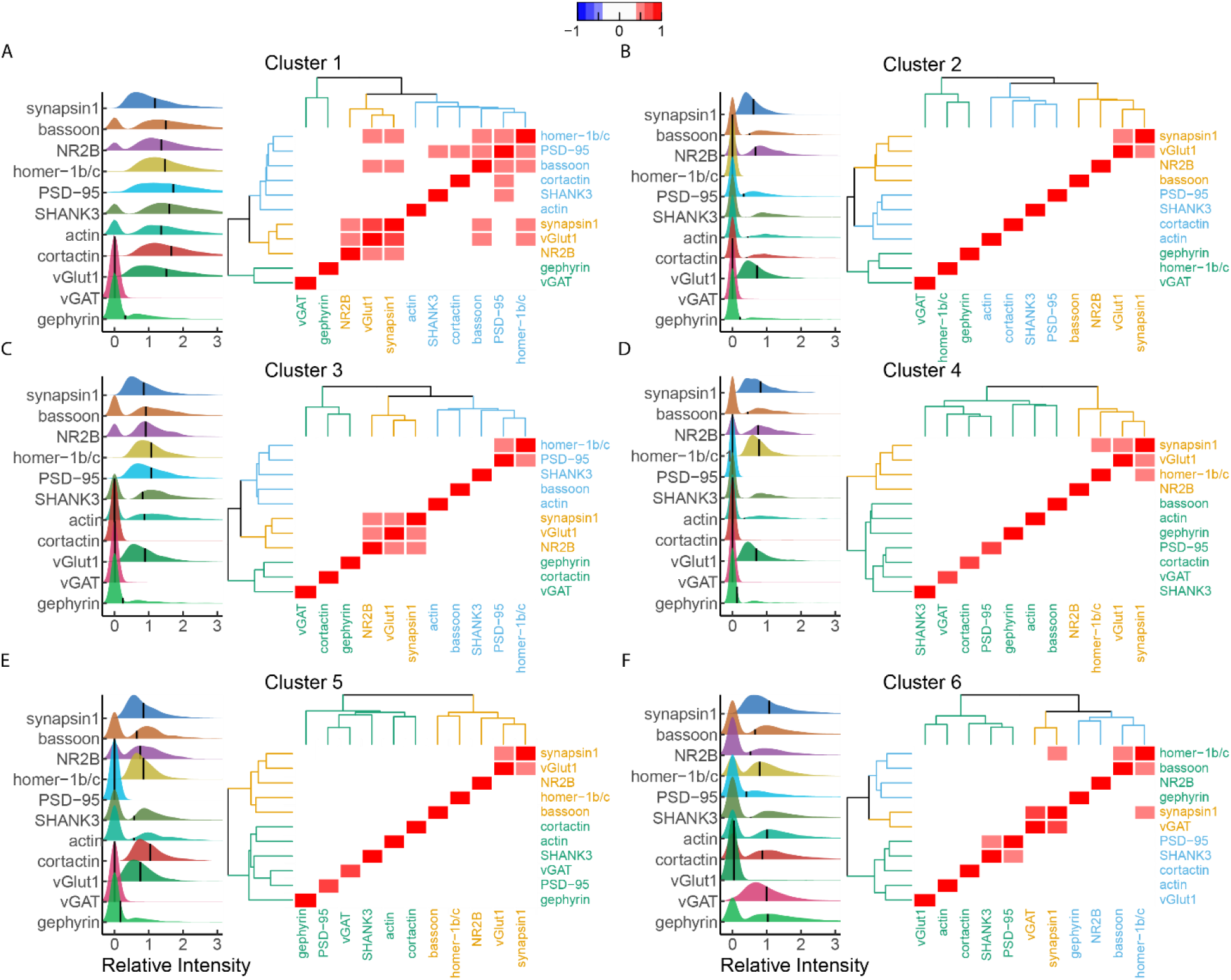
Comparison of synaptic intensity and protein relationships among all proteins within each cluster. Ridgeline plots of relative synaptic intensity for each cluster group identified using HDBScan. Horizontal black line represents cluster mean intensity. All values are normalized to untreated mean integrated intensity. Heatmap indicates the correlation coefficient between each protein. Correlation values between −0.4 and 0.4 obscured to highlight strong correlations. Dendrograms surrounding heatmap show hierarchical clustering of proteins within each cluster.

Ridgeline plots (**Figure 3,** left subpanel) were created for each protein within each cluster to compare the relative synaptic levels of each target and to examine the proportion of synapses with detectable (> 0) levels of each target. Additionally, pairwise correlation analysis of the synaptic protein levels was performed to identify proteins with coordinated synaptic expression (**Figure 3,** right subpanel). Hierarchical clustering was then performed on the correlation matrix to identify which groups of proteins had similar levels of coordinated synaptic expression. Comparing these features across the excitatory clusters shows that Cluster 1 had the highest level of synaptic proteins, the lowest proportion of synapses with undetectable levels of synaptic proteins (**Figure 3A-E.** left subpanel), and the most proteins with coordinated synaptic expression (**Figure 3A-E**, right subpanel) of the excitatory synaptic clusters. Cluster 3 was the most similar to Cluster 1 as it had the next highest synaptic levels of proteins and the next highest number of proteins with coordinated synaptic expression. Clusters 2, 4 and 5 had lower levels of proteins at synapses compared to Cluster 1 and 3, more synapses with undetectable levels of proteins, and very few proteins with coordinated synaptic expression. In these clusters the proteins with most of the coordinated synaptic expression were synapsin1 and vGlut1. Taken together, these clusters may be synapses in various stages of maturation, synapses actively remodeling or they could represent false synapses caused from staining artifacts.

Cluster 6 contains inhibitory synapses with vGAT detectable in 97.8% of these synapses. Gephyrin levels were also the highest within this cluster relative to all other clusters, but, interestingly, were only detectable in 61% of these synapses. The excitatory postsynaptic scaffolds SHANK3 and PSD-95 were assigned to inhibitory synapses in ~41% and 36% of the synapses in this cluster. Additionally, homer-1b/c was also found near inhibitory synapses in 73% of these synapses but with lower intensity with respect to Cluster 1 (large effect > 0.8) (**Figure 3 and Supplemental Figure 17**). Correlation analysis of proteins within this cluster revealed coordinated synaptic expression between vGAT and synapsin1, between PSD-95 and SHANK3, between homer-1b/c and bassoon and surprisingly between synapsin1 and homer-1b/c. Taken together, within this cluster we only see coordinate synaptic expression among the presynaptic inhibitory synaptic proteins. Additionally, our results suggest homer-1b/c may serve some yet unknown function at inhibitory synapses.

### Response of excitatory and inhibitory synapses to activity blockade

The phenomena of neurons eliciting compensatory structural changes within synapses to maintain network homeostatic balance in response to chronic activity blockade has been characterized in numerous studies (G. G. Turrigiano 2008). However, the overview of simultaneous changes in inhibitory and excitatory synapses has been limited by conventional immunocytochemistry. We used PRISM staining to simultaneously investigate the effect of chronic activity blockade on excitatory and inhibitory synapses. For this, the distribution pattern of excitatory and inhibitory synapses was determined by using the classification method established above. We found a higher abundance of excitatory inputs compared to inhibitory on cultured neurons (**Figure 4B**). Under control conditions, glutamatergic synapses have a density of 56±9.6 synapses per 100 μm length of dendrite, whereas GABAergic synapses density was 15.8±9.5 per 100 μm; which gives an average excitatory to inhibitory synapses ratio of 5+3:1 calculated per replicate (**Figure 4B**). Accordingly with the well-established model that neurons coordinate their excitatory and inhibitory inputs to establish and maintain a constant excitation/inhibition (E/I) ratio which is essential for circuit function and stability (He and Cline 2019; Zhou et al. 2014; He et al. 2018), we observed that TTX treatment did not alter this ratio or density of synapses in hippocampal cultured neurons (**Figure 4A and B**).

**Figure 4.**
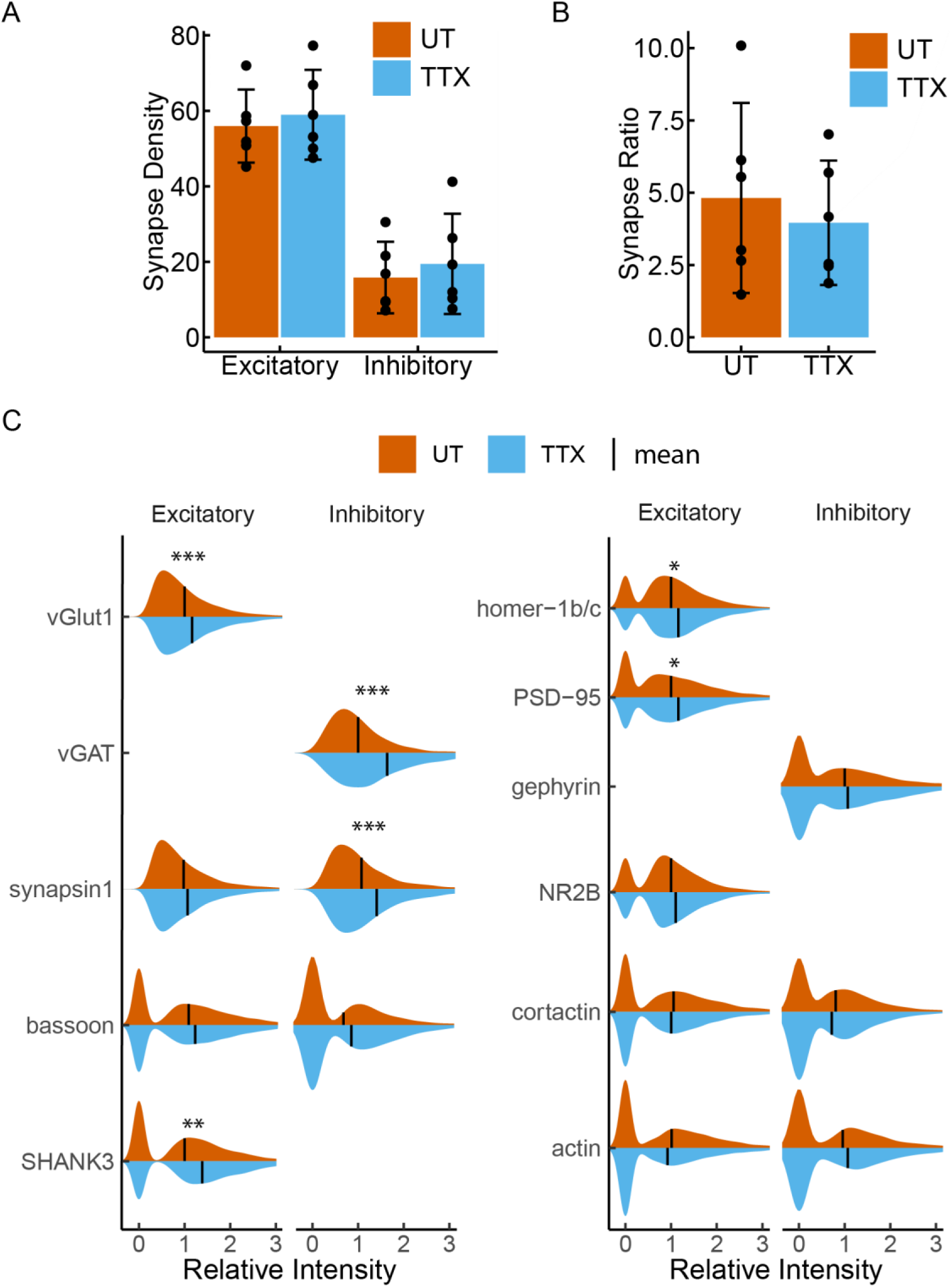
Excitatory and inhibitory synaptic density does not change in response to 48hr TTX treatment. (A) Quantification of excitatory and inhibitory synaptic density from untreated (red) and TTX treated (blue) cells. Bar height represents the mean number of synapses per 100 μm length of dendrite. (B) Quantification of the excitatory: inhibitory synaptic ratio in untreated (red) and TTX treated (blue) neurons. Bar height represents mean excitatory:inhibitory ratio. Error bars indicate 95% confidence intervals. Closed circles indicate results from individual replicates n = 6. (C) Violin plots of relative synaptic intensity for synaptic targets at excitatory synapses (left) and inhibitory synapses (right) from untreated (red) and TTX treated cells (blue). Black line indicates the mean intensity. P-values are computed using Student’s t-test on the mean Integrated Intensity with n = 6 (UT) and n = 5 (TTX) replicates. All values are normalized to untreated mean integrated intensity. *** indicates p < 0.001; ** indicates p < 0.01; * indicates p < 0.05.

It is well known that homeostatic plasticity regulates the relative strength of excitatory and inhibitory synapses in order to keep relatively stable firing rates of neurons by increasing synaptic levels of proteins that regulate excitatory signaling (G. G. Turrigiano et al. 1998; G. G. Turrigiano and Nelson 2000; G. G. Turrigiano 2008; G. G. Turrigiano and Nelson 2004; G. Turrigiano 2012; Liu 2004; Hartman et al. 2006). We applied our imaging platform and CellProfiler software to identify excitatory and inhibitory synapses and to examine the response of 11 synaptic targets simultaneously following induction of homeostatic synaptic plasticity. Interestingly, using relative intensity measurements to evaluate protein abundance, we observed different patterns of response to neuronal activity blockade depending whether proteins are present at excitatory or inhibitory synapses or are pre-or postsynaptic (**Figure 4C**). The neurotransmitter transporters at presynaptic excitatory and inhibitory terminals, vGlut1 and vGAT, respectively, both show a significant increase in response to TTX treatment. In contrast, synapsin1, a protein associated with the reserve pool of synaptic vesicles, shows increased levels in response to TTX treatment exclusively at inhibitory synapses. Additionally, at both excitatory and inhibitory synapses there was no change in the levels of the presynaptic protein bassoon. However, overall bassoon levels were lower at inhibitory synapses compared to excitatory synapses. On the postsynaptic side, the levels of the excitatory scaffolding proteins SHANK3, homer-1b/c, and PSD-95 increased in the presence of TTX, while no changes in the inhibitory postsynaptic scaffold gephyrin were observed at inhibitory synapses. Interestingly, no changes were observed in the cytoskeletal proteins cortactin and actin at excitatory or inhibitory synapses (**Figure 4C**). Thus, we observe that excitatory synapses respond both postsynaptically and presynaptically to neuronal activity suppression while at inhibitory synapses the response is mainly presynaptic.

### Alterations in excitatory and inhibitory synapses within synaptic subgroups following activity blockade

We further assessed the effect of activity blockage via TTX on synaptic sub-types and found that the same clusters described in control neurons are present after chronic neuronal activity blockage (**Figure 5**). Power analysis of synaptic intensity revealed negligible differences (effect < 0.2) for most synaptic targets within each cluster (**Figure 5B**). Instead we found that following 48-hr TTX treatment, the number of synapses present in each cluster changed. Specifically, there was a small decrease in the number of synapses within Clusters 2, 4, and 5 (p = 0.05, p = 0.13, and p < 0.001) and a small increase in the number of synapses in the Clusters 1 and 3 (p = 0.09 and p = 0.08, respectively) (**Figure 5C**). Taken together the results indicate that TTX treatment does not produce unique synaptic subtypes (based on the proteins we examined) but it does change the number of synapses present within each cluster, specifically, reducing the number of synapses in clusters defined by low protein expression and increasing the number of synapses in clusters defined by higher protein expression.

**Figure 5:**
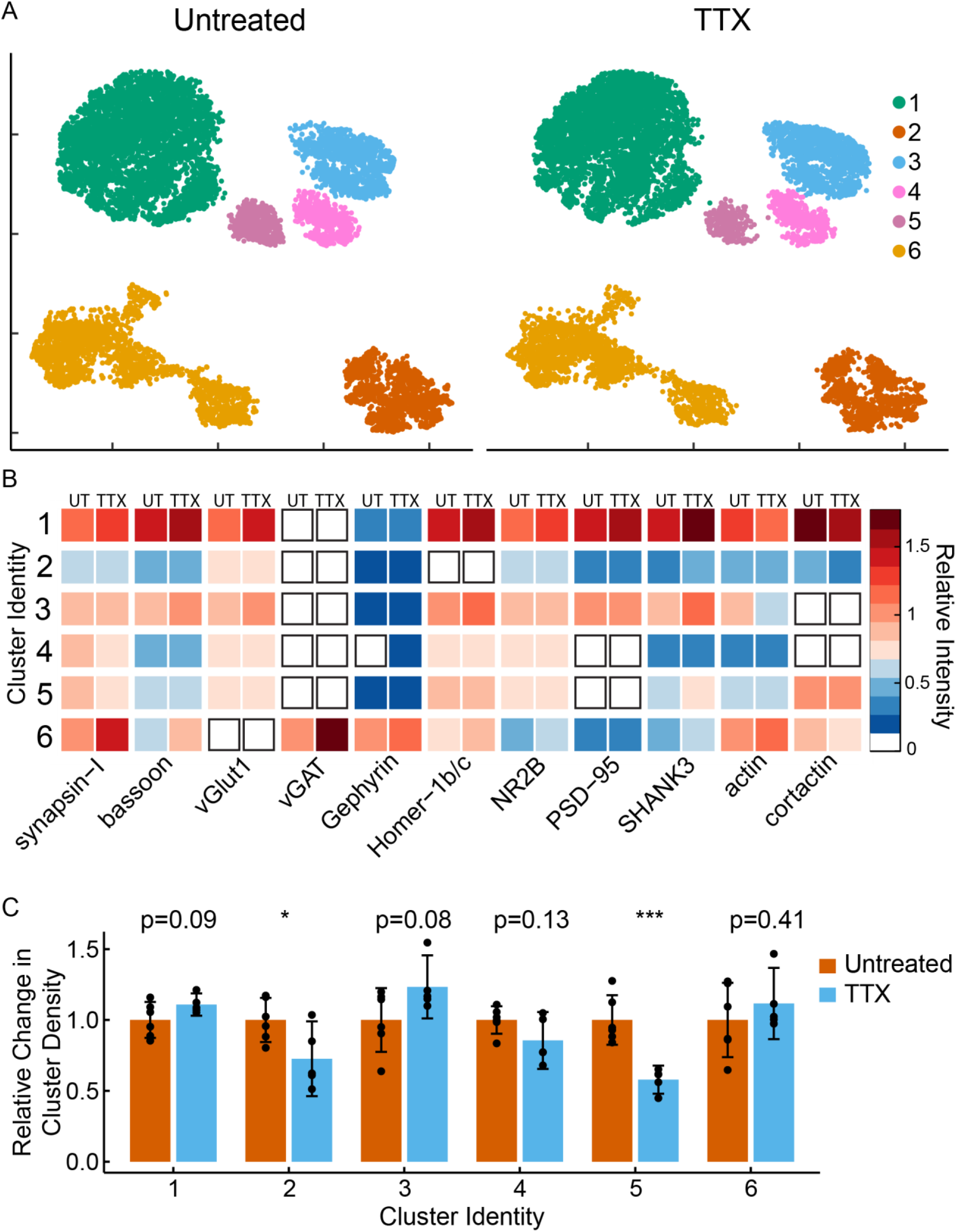
Characterizing changes within clusters in response to TTX treatment. (A) UMAP plots of individual synapses (n = 10,000) separated by treatment group. (B) Heatmap indicates the average relative intensities for each synaptic target within each cluster. Left side of the column is the untreated sample, the right side of the column is the TTX treated sample. All values are normalized to untreated mean integrated intensity. (C) Bar heights show synapse density, relative to untreated groups, within each HDBSCAN identified cluster following TTX treatment. P-values are computed using Student’s t-test with n = 6 (UT) and n = 5 (TTX) replicates; p < 0.001 (***).

## Discussion

We simultaneously characterized inhibitory and excitatory synaptic content from multiplexed imaging using PRISM and CellProfiler, offering detailed molecular insight into synapse complexity at the single-synapse level, including excitatory and inhibitory components.

Synapse identification using synapsin1 in combination with either vGlut1 or vGAT indicated an excitatory-to-inhibitory (E/I) ratio of approximately 5±3:1 in hippocampal neurons. In cultured neurons, the E/I ratio reported using imaging techniques is variable and dependent on multiple factors such as brain region and the age of the culture. In hippocampal culture, E/I ratios range between 2.5:1 to 17:1 (Harrill et al. 2015; Liu 2004; Gulyás et al. 1999; Megías et al. 2001), which may be a consequence of the synaptic distribution variations of across the dendrites with fewer synapses located proximal than distal segments (Megías et al. 2001; Bloss et al. 2016). In agreement with this, the methodology implemented here provides the overall E/I ratios from whole synapse populations based on the use of several protein markers simultaneously.

Beyond E/I ratios, simultaneous imaging of 11 synaptic components enabled the identification and labeling of synapses based on synaptic content, and characterization of unique changes within these synapse sub-populations in response to chemical perturbation. Considering only those synapses that met the classification criteria for excitatory and inhibitory described here, we were able to identify 5 types of excitatory synapses and 1 type of inhibitory synapse based on the differential expression patterns of the targets specifically selected for this study. These synaptic targets, which are all major components of synapses, have been extensively studied for their role in synapse structure and function (Morgan Sheng and Hoogenraad 2007; M. Sheng and Kim 2011). Although the significance of these clusters may require additional studies due to the disproportionate complexity of the synaptic proteome (Husi et al. 2000; Peng et al. 2004; Collins et al. 2005; 2006; Richard David Emes and Grant 2011; Distler et al. 2014). Notwithstanding, our analysis offers a framework for deciphering orchestrated compensatory changes that occur during synaptic scaling.

An additional layer of complexity that requires consideration, when classifying synapses using protein levels and correlation analysis, is the highly dynamic and plastic nature of synapses (Nimchinsky, Sabatini, and Svoboda 2002). Given this framework, the clusters we identified may represent synapses captured in various stages of remodeling, which may explain the diversity observed, especially regarding excitatory synapse subtypes. Another factor to consider is the influence of developmental factors on the morphological, structural, and proteomic characteristics of synapses. Thus, an alternative interpretation of these clusters is that we captured synapses from various developmental stages.

We must also acknowledge the possibility that, due to the variation of antibody epitope specificity among different host species and purification methods, some of the identified clusters may indicate contamination from staining artifacts. While visual inspection of these synapses does not present any evidence to suggest this is the case, assessing detection of false positives could serve as a useful tool for improving accuracy.

Another interesting observation from our results is that excitatory synapses seem to be more heterogeneous than inhibitory synapses. Excitatory synapses present more diverse morphologies, which could account for the increased variation. However, we only incorporated two inhibitory markers, vGAT and gephyrin, which would limit the number of inhibitory clusters that could be detected. While these interpretations represent exciting possibilities for examination of synapse biology, further experimentation to test these hypotheses are critical to address all these different aspects associated with synapse diversity classification.

Another unexpected advantage of the platform implemented here is that it also provides a tool for the simultaneous examination of the population of dually innervated synapses, originally described by electron microscopy (Kubota et al. 2016; Villa et al. 2016). This population, observed here as positive for both vGlut1 and vGAT(12%), will likely require detailed characterization using super-resolution microscopy to overcome the resolution limitations of our confocal microscope(187 nm). Thus, this platform could be used in tandem with DNA-PRISM, which also has super-resolution capabilities (Guo et al. 2019).

Our results revealed differences in the responses to activity blockade at excitatory and inhibitory synapses which was not described previously (Guo et al. 2019). At excitatory synapses, the synaptic levels of postsynaptic scaffolding proteins and presynaptic neurotransmitter transporter levels increased whereas there were no changes in bassoon levels or changes in the presynaptic vesicle clustering protein synapsin1. Our results suggest that, at excitatory synapses, neurons respond to activity blockade by increasing the amount of neurotransmitter present per vesicle and increasing the available postsynaptic AMPA receptor binding slots, but not by increasing the number of synaptic vesicles that are available for release. In contrast, inhibitory synapses showed both an increase in the neurotransmitters amount in the vesicle and an increase in the reserve pool of vesicles, indicated by synapsin1. Synapsin1 has a critical function associated with the reserve pool of synaptic vesicles (Bloom et al. 2003; Akbergenova and Bykhovskaia 2007; Gitler et al. 2008; Hilfiker et al. 1998), regulating mobilization of vesicles into the recycling pool (Chi, Greengard, and Ryan 2003; Cousin et al. 2003; Menegon et al. 2006; Baldelli et al. 2007) and forming clusters of vesicles that are reluctant to release unless high frequency stimulation is applied (Rizzoli and Betz 2005). Increased synapsin1 at inhibitory synapses could therefore indicate reduced baseline GABAergic signaling consistent with the conventional model of synaptic scaling.

Synaptic diversity across various brain regions has been described previously using two postsynaptic markers PSD-95 and SAP102 (Zhu et al. 2018) however, a detailed characterization of synaptic diversity in cultured neurons and their response to changes in network activity with high-content imaging has not been addressed. An important component of homeostatic plasticity is that the resultant changes in excitability maintain the relative differences in synaptic strength between the individual synapses. Although no differences in the synapse diversity were observed in the presence of TTX treatment compared to control conditions; TTX was able to modify the clusters producing fewer synapses defined by low protein levels, consistent with the model that synaptic scaling induces global changes to increase synaptic strength. While this was true for synapses defined by our synaptic targets, there may be additional variations that require different targets for detection. Additionally, our observations of synaptic diversity are from hippocampal neurons grown in culture which is a useful tool for studies of synapse development and plasticity but lacks the organization level and experience driven plasticity present *in vivo* models. Incorporation of additional targets and application to different model systems should reveal exciting discoveries.

In conclusion, PRISM facilitated the exploration of complex synaptic architecture, with the imaging and analysis platform described here revealing numerous synaptic subtypes and their molecular rearrangements in response to homeostatic scaling. Exploration of synaptic diversity in neurons enables a new range of investigations with the potential to reveal new connections between synaptic architecture and neuronal function that can be especially useful for high-content and high throughput screening using compounds or siRNA libraries of genes-associated with disease.

## Methods

### Primary mouse and rat neuronal cultures

Procedures for rat neuronal culture were reviewed and approved for use by the Broad Institutional Animal Care and Use Committee. For rat hippocampal neuronal cultures, E18 embryos were collected from CO_2_ euthanized pregnant Sprague Dawley rats (Taconic). Embryo hippocampi were dissected in 4 °C Hibernate E supplemented with 2% B27 supplements and 100 U/mL Penicillin/Strep (Thermo Fisher Scientific). Hippocampal tissues were digested in Hibernate E containing 20 U/mL papain, 1 mM L-cysteine, 0.5 mM EDTA (Worthington Biochem) and 0.01% DNAse (Sigma-Aldrich) for 8 min, and was stopped with 0.5% ovomucoid trypsin inhibitor (Worthington Biochem) and 0.5% bovine serum albumin (BSA) (Sigma-Aldrich). Neurons were plated at a density of 15,000 cells/well onto poly-D-lysine coated, black-walled, thin-bottomed 96-well plates (Greiner Bio-One). Neurons were seeded and maintained in NbActiv1(BrainBits). Cells were grown at 37 °C in 95% air with a 5% CO_2_ humidified incubator for 19 days before use. Cells were treated on day 19 with 1 μM TTX (Sigma-Aldrich) for 48 hrs. All procedures involving animals were in accordance with the US National Institutes of Health Guide for the Care and Use of Laboratory Animals.

### Immunostaining for LNA-PRISM

Cells were fixed in 4% paraformaldehyde in and 4% wt/vol sucrose in PBS for 15 min at room temperature and permeabilized with 0.25% Triton X100 in PBS. Permeabilized cells were incubated in 50 μg/ml RNase A (Thermo Fisher Scientific) and 230 U/mL RNase T1 (Thermo Fisher Scientific) in PBS at 37°C for 1 h to reduce the fluorescent background caused by ssLNA-RNA binding, and washed 3 times with PBS. Cells were then blocked for 1 hr at room temperature with the 5% BSA in PBS. The first round of primary staining was performed using unconjugated primary antibodies diluted in the regular blocking buffer: MAP2, VGLUT1, PSD-95, NR2B, and gephyrin. Cells were incubated in primary antibodies overnight at 4 °C, washed 3 times with PBS, and then incubated in the nuclear blocking buffer for 1 hr at room temperature. The first round of secondary staining antibodies were diluted in the nuclear blocking buffer and incubated for 1 hr at room temperature. Antibodies used in the first round of secondary are: goat-anti-chicken-Alexa 488, goat-anti-guinea pig-Alexa555 and goat-anti-rabbit-p1, goat-anti-mouse-p12, goat-anti-rat-p7. Cells were washed 3 times with PBS, post-fixed for 15 min with 4% paraformaldehyde and 4% wt/vol sucrose in PBS. Cells were washed 3 times with PBS and incubated again in the nuclear blocking buffer for 30 min at room temperature. The second round of primary antibodies were incubated overnight at 4 °C in the following primary antibody/peptide solution diluted in the nuclear blocking buffer using: phalloidin-p2, vGAT-p3, cortactin-p4, SHANK3-p6, bassoon-p8, synapsin1-p9, homer-1b/c-p10. Cells were then washed 3 times with PBS then incubated in diluted DAPI for 15 min. See **Supplementary Table 2** for detailed antibody information and antibody conjugation method.

### Multiplexed confocal imaging of neurons using LNA-PRISM

LNA-PRISM imaging was performed using the Opera Phenix High-Content Screening System (PerkinElmer) equipped with a fast laser-based autofocus system, high NA water immersion objective (63x, numerical aperture = 1.15), two large format scientific complementary metal-oxide semiconductor (sCMOS) cameras and spinning disk optics. 405 nm, 488 nm and 561 nm lasers were used as excitation for DAPI, MAP2, and vGlut1 channels respectively. PRISM images were acquired using a 640 nm laser (40 mW), sCMOS camera with 1-2 s exposure time, and effective pixel size of 187 nm. Imaging probe was freshly diluted to 10 nM in imaging buffer (500 mM NaCl in PBS, pH 8) immediately before imaging. Neurons were incubated with imaging probes for 5 min and washed twice with imaging buffer to remove unbound probe. See **Supplemental Table 3** for LNA imager probe sequences. For each field of view, a stack of three images was acquired with a step of 0.5 μm. At least five fields of view were imaged for each well. Following imaging, cells were washed two times with wash buffer (0.01 x PBS) for 3 minutes each round. To account for variabilities within each plate and across different cultures, imaging was performed on two independent neuronal cultures with three wells imaged from each neuronal culture.

### Image analysis

CellProfiler (McQuin et al. 2018) is a popular image analysis tool containing numerous image segmentation methods and analysis tools. The pipeline was modeled after our original method (Guo et al. 2019). Several minor modifications to the original procedure were implemented to improve image segmentation quality and to make our method compatible with CellProfiler (**Supplementary Table 5**). The pipeline can be divided into three main stages. In the first stage the images are imported, aligned and uneven illumination is corrected. The output from this first stage of the pipeline is the aligned tiff images with the corrected background. These images serve as a quality control checkpoint to ensure correct image alignment and illumination correction. The second stage of the pipeline performs image segmentation on the images to define and locate the nuclei, dendrites and puncta from each round of imaging. The pipeline offers users the ability to easily customize key aspects of the analysis methodology such as illumination correction and thresholding without any source code modifications. This stage of the pipeline produces binary images of the object segmentation and also the grey-scale images of the puncta channels following application of the white-top hat filters used to enhance puncta segmentation. These images serve as helpful guides to ensure optimal image segmentation. The final stage of the pipeline groups all of the segmented objects into synapses, based on the level of overlap with the synapsin1 objects, nuclear masks and the dendritic masks. For our analysis puncta were considered non-synaptic if the puncta overlapped the nuclei region, were outside of the dendrite object masks or did not overlap with synapsin1. Postsynaptic objects with at least 6.25% overlap with synapsin1 were considered synaptic. Presynaptic objects were considered synaptic if the overlap was at least 50%. The final stage of the pipeline outputs binary images of the synaptic objects and multiple csv files containing the quantification of the synaptic objects. An additional modification from our original method was the incorporation of two inhibitory synaptic markers: vGAT and gephyrin. Synapses were defined as excitatory when only vGlut1 puncta were present and as inhibitory when only vGAT puncta were present. For quality control of clustering we isolated and plotted synapses from individual wells separately to ensure identified clusters were not artifacts resulting from differences in staining or image intensity among the image sets (**Supplemental Figure 18**). The pipeline is available as **Supplemental File 1.**

### Statistical analysis

All statistical analysis was performed using R statistical environment. For the ridgeline plots 2000 synapses were randomly picked from each replicate to create the final plot, to ensure each replicate contributed equally to the final result. A two-tailed Students’ T-Test of the mean Integrated Intensity was used to determine p-value. Plots created using ggridges package in R (Wilke 2018). For bar-chart comparison a two-tailed Student’s t-test was used to determine a significant (p < 0.05) difference between groups. Barplots created using ggplot2 and ggpubr packages in R (Wickham 2016; Kassambara 2019). UMAP was performed using the umap package in R stats (Konopka 2019). For UMAP clustering 2000 synapses were randomly subsampled from each well. Columns containing missing data (NA) were removed. Feature reduction was performed using findCorrelation in caret package of R stats (Wing et al. 2019). Remaining data was centered using the scale function of R Stats before using the data as input for UMAP. The resulting UMAP object was used to transform all remaining data. HDBScan was performed on umap output using hdbscan module from python with min_cluster_size = 100 (McInnes, Healy, and Astels 2017). Data considered as noise from HDBscan was removed from the plot. Scatterplots were generated using ggplot2 and ggpubr. Heatmaps were created using gplots R package (Warnes et al. 2020).

Individual correlation matrices were generated for each biological replicate within each UMAP group and then averaged to produce a final representative correlation matrix for each UMAP group. Correlation values between −0.4 and 0.4 obscured to highlight strong correlations. Effect size was calculated using integrated intensity measurements from individual synapses, n = 2000 for each replicate. Effect size calculated using effsize R package (Torchiano 2016).

## Acknowledgements

Funding from the NIH R01-MH112694 to M.B. and J.R.C. and the NSF Physics of Living Systems 1707999 to M.B. is gratefully acknowledged. J.R.C. acknowledges funding from the Stanley Center for Psychiatric Research.

## Author contributions

E.D., K.P., M.B., and J.R.C conceived of and designed the experimental study. K.P. prepared the primary neuronal cultures and performed the TTX treatments. E.W. prepared the PRISM imaging reagents. K.P. and E.D. performed PRISM imaging. B.C. designed and implemented the CellProfiler image analysis pipeline. E.D., K.P., B.C. and S.S. analyzed and interpreted the PRISM data. M.B., A.E.C., and J.R.C. supervised the overall study. All authors wrote and commented on the manuscript.

## References

Akbergenova, Yulia, and Maria Bykhovskaia. 2007. “Synapsin Maintains the Reserve Vesicle Pool and Spatial Segregation of the Recycling Pool in Drosophila Presynaptic Boutons.” Brain Research 1178 (October): 52–64. https://doi.org/10.1016/j.brainres.2007.08.042.

Baldelli, P., A. Fassio, F. Valtorta, and F. Benfenati. 2007. “Lack of Synapsin I Reduces the Readily Releasable Pool of Synaptic Vesicles at Central Inhibitory Synapses.” Journal of Neuroscience 27 (49): 13520–31. https://doi.org/10.1523/JNEUROSCI.3151-07.2007.

Bayés, Àlex, Mark O. Collins, Mike D. R. Croning, Louie N. van de Lagemaat, Jyoti S. Choudhary, and Seth G. N. Grant. 2012. “Comparative Study of Human and Mouse Postsynaptic Proteomes Finds High Compositional Conservation and Abundance Differences for Key Synaptic Proteins.” Edited by Anna Dunaevsky. PLoS ONE 7 (10): e46683. https://doi.org/10.1371/journal.pone.0046683.

Bayés, Àlex, Mark O. Collins, Rita Reig-Viader, Gemma Gou, David Goulding, Abril Izquierdo, Jyoti S. Choudhary, Richard D. Emes, and Seth G. N. Grant. 2017. “Evolution of Complexity in the Zebrafish Synapse Proteome.” Nature Communications 8 (1): 14613. https://doi.org/10.1038/ncomms14613.

Bayés, Àlex, Louie N van de Lagemaat, Mark O Collins, Mike D R Croning, Ian R Whittle, Jyoti S Choudhary, and Seth G N Grant. 2011. “Characterization of the Proteome, Diseases and Evolution of the Human Postsynaptic Density.” Nature Neuroscience 14 (1): 19–21. https://doi.org/10.1038/nn.2719.

Bloom, Ona, Emma Evergren, Nikolay Tomilin, Ole Kjaerulff, Peter Löw, Lennart Brodin, Vincent A. Pieribone, Paul Greengard, and Oleg Shupliakov. 2003. “Colocalization of Synapsin and Actin during Synaptic Vesicle Recycling.” The Journal of Cell Biology 161 (4): 737–47. https://doi.org/10.1083/jcb.200212140.

Bloss, Erik B., Mark S. Cembrowski, Bill Karsh, Jennifer Colonell, Richard D. Fetter, and Nelson Spruston. 2016. “Structured Dendritic Inhibition Supports Branch-Selective Integration in CA1 Pyramidal Cells.” Neuron 89 (5): 1016–30. https://doi.org/10.1016/j.neuron.2016.01.029.

Chi, Ping, Paul Greengard, and Timothy A Ryan. 2003. “Synaptic Vesicle Mobilization Is Regulated by Distinct Synapsin I Phosphorylation Pathways at Different Frequencies.” Neuron 38 (1): 69–78. https://doi.org/10.1016/S0896.6273(03)00151-X.

Collins, Mark O., Holger Husi, Lu Yu, Julia M. Brandon, Chris N. G. Anderson, Walter P. Blackstock, Jyoti S. Choudhary, and Seth G. N. Grant. 2006. “Molecular Characterization and Comparison of the Components and Multiprotein Complexes in the Postsynaptic Proteome.” Journal of Neurochemistry 97 (April): 16–23. https://doi.org/10.1111/j.1471-4159.2005.03507.x.

Collins, Mark O., Lu Yu, Marcelo P. Coba, Holger Husi, Iain Campuzano, Walter P. Blackstock, Jyoti S. Choudhary, and Seth G. N. Grant. 2005. “Proteomic Analysis of *in Vivo* Phosphorylated Synaptic Proteins.” Journal of Biological Chemistry 280 (7): 5972–82. https://doi.org/10.1074/jbc.M411220200.

Cousin, Michael A., Chandra S. Malladi, Timothy C. Tan, Clarke R. Raymond, Karen J. Smillie, and Phillip J. Robinson. 2003. “Synapsin I-Associated Phosphatidylinositol 3-Kinase Mediates Synaptic Vesicle Delivery to the Readily Releasable Pool.” Journal of Biological Chemistry 278 (31): 29065–71. https://doi.org/10.1074/jbc.M302386200.

Distler, Ute, Michael J. Schmeisser, Assunta Pelosi, Dominik Reim, Jörg Kuharev, Roland Weiczner, Jan Baumgart, et al. 2014. “In-Depth Protein Profiling of the Postsynaptic Density from Mouse Hippocampus Using Data-Independent Acquisition Proteomics.” PROTEOMICS 14 (21–22): 2607–13. https://doi.org/10.1002/pmic.201300520.

Emes, Richard D., and Seth G.N. Grant. 2012. “Evolution of Synapse Complexity and Diversity.” Annual Review of Neuroscience 35 (1): 111–31. https://doi.org/10.1146/annurev-neuro-062111-150433.

Emes, Richard D, Andrew J Pocklington, Christopher N G Anderson, Alex Bayes, Mark O Collins, Catherine A Vickers, Mike D R Croning, et al. 2008. “Evolutionary Expansion and Anatomical Specialization of Synapse Proteome Complexity.” Nature Neuroscience 11 (7): 799–806. https://doi.org/10.1038/nn.2135.

Emes, Richard David, and Seth G. N. Grant. 2011. “The Human Postsynaptic Density Shares Conserved Elements with Proteomes of Unicellular Eukaryotes and Prokaryotes.” Frontiers in Neuroscience 5. https://doi.org/10.3389/fnins.2011.00044.

Gerdes, M. J., C. J. Sevinsky, A. Sood, S. Adak, M. O. Bello, A. Bordwell, A. Can, et al. 2013. “Highly Multiplexed Single-Cell Analysis of Formalin-Fixed, Paraffin-Embedded Cancer Tissue.” Proceedings of the National Academy of Sciences 110 (29): 11982–87. https://doi.org/10.1073/pnas.1300136110.

Gitler, D., Q. Cheng, P. Greengard, and G. J. Augustine. 2008. “Synapsin IIa Controls the Reserve Pool of Glutamatergic Synaptic Vesicles.” Journal of Neuroscience 28 (43): 10835–43. https://doi.org/10.1523/JNEUROSCI.0924-08.2008.

Gulyás, A. I., M. Megías, Z. Emri, and T. F. Freund. 1999. “Total Number and Ratio of Excitatory and Inhibitory Synapses Converging onto Single Interneurons of Different Types in the CA1 Area of the Rat Hippocampus.” The Journal of Neuroscience: The Official Journal of the Society for Neuroscience 19 (22): 10082–97.

Guo, Syuan-Ming, Remi Veneziano, Simon Gordonov, Li Li, Eric Danielson, Karen Perez de Arce, Demian Park, et al. 2019. “Multiplexed and High-Throughput Neuronal Fluorescence Imaging with Diffusible Probes.” Nature Communications 10 (1): 4377. https://doi.org/10.1038/s41467-019-12372-6.

Harrill, Joshua A., Hao Chen, Karin M. Streifel, Dongren Yang, William R. Mundy, and Pamela J. Lein. 2015. “Ontogeny of Biochemical, Morphological and Functional Parameters of Synaptogenesis in Primary Cultures of Rat Hippocampal and Cortical Neurons.” Molecular Brain 8 (February): 10. https://doi.org/10.1186/s13041-015-0099-9.

Hartman, Kenichi N, Sumon K Pal, Juan Burrone, and Venkatesh N Murthy. 2006. “Activity-Dependent Regulation of Inhibitory Synaptic Transmission in Hippocampal Neurons.” Nature Neuroscience 9 (5): 642–49. https://doi.org/10.1038/nn1677.

He, Hai-yan, and Hollis T Cline. 2019. “What Is Excitation/Inhibition and How Is It Regulated? A Case of the Elephant and the Wisemen.” Journal of Experimental Neuroscience 13 (January): 117906951985937. https://doi.org/10.1177/1179069519859371.

He, Hai-yan, Wanhua Shen, Lijun Zheng, Xia Guo, and Hollis T. Cline. 2018. “Excitatory Synaptic Dysfunction Cell-Autonomously Decreases Inhibitory Inputs and Disrupts Structural and Functional Plasticity.” Nature Communications 9 (1): 2893. https://doi.org/10.1038/s41467-018-05125-4.

Hilfiker, Sabine, Felix E. Schweizer, Hung-Teh Kao, Andrew J. Czernik, Paul Greengard, and George J. Augustine. 1998. “Two Sites of Action for Synapsin Domain E in Regulating Neurotransmitter Release.” Nature Neuroscience 1 (1): 29–35. https://doi.org/10.1038/229.

Husi, Holger, Malcolm A. Ward, Jyoti S. Choudhary, Walter P. Blackstock, and Seth G. N. Grant. 2000. “Proteomic Analysis of NMDA Receptor-Adhesion Protein Signaling Complexes.” Nature Neuroscience 3 (7): 661–69. https://doi.org/10.1038/76615.

Kassambara, Alboukadel. 2019. Ggpubr: “ggplot2” Based Publication Ready Plots. https://CRAN.R-project.org/package=ggpubr.

Konopka, Tomasz. 2019. Umap: Uniform Manifold Approximation and Projection. https://CRAN.R-project.org/package=umap.

Kubota, Yoshiyuki, Fuyuki Karube, Masaki Nomura, and Yasuo Kawaguchi. 2016. “The Diversity of Cortical Inhibitory Synapses.” Frontiers in Neural Circuits 10 (April). https://doi.org/10.3389/fncir.2016.00027.

Lin, Jia-Ren, Mohammad Fallahi-Sichani, and Peter K. Sorger. 2015. “Highly Multiplexed Imaging of Single Cells Using a High-Throughput Cyclic Immunofluorescence Method.” Nature Communications 6 (1): 8390. https://doi.org/10.1038/ncomms9390.

Liu, Guosong. 2004. “Local Structural Balance and Functional Interaction of Excitatory and Inhibitory Synapses in Hippocampal Dendrites.” Nature Neuroscience 7 (4): 373–79. https://doi.org/10.1038/nn1206.

Mclnnes, Leland, John Healy, and Steve Astels. 2017. “Hdbscan: Hierarchical Density Based Clustering.” The Journal of Open Source Software 2 (11). https://doi.org/10.21105/joss.00205.

McQuin, Claire, Allen Goodman, Vasiliy Chernyshev, Lee Kamentsky, Beth A. Cimini, Kyle W. Karhohs, Minh Doan, et al. 2018. “CellProfiler 3.0: Next-Generation Image Processing for Biology.” Edited by Tom Misteli. PLOS Biology 16 (7): e2005970. https://doi.org/10.1371/journal.pbio.2005970.

Megías, M, Zs Emri, T.F Freund, and A.I Gulyás. 2001. “Total Number and Distribution of Inhibitory and Excitatory Synapses on Hippocampal CA1 Pyramidal Cells.” Neuroscience 102 (3): 527–40. https://doi.org/10.1016/S0306-4522(00)00496-6.

Menegon, A., D. Bonanomi, C. Albertinazzi, F. Lotti, G. Ferrari, H.-T. Kao, F. Benfenati, P. Baldelli, and F. Valtorta. 2006. “Protein Kinase A-Mediated Synapsin I Phosphorylation Is a Central Modulator of Ca2+-Dependent Synaptic Activity.” Journal of Neuroscience 26 (45): 11670–81. https://doi.org/10.1523/JNEUROSCI.3321-06.2006.

Nimchinsky, Esther A., Bernardo L. Sabatini, and Karel Svoboda. 2002. “Structure and Function of Dendritic Spines.” Annual Review of Physiology 64 (1): 313–53. https://doi.org/10.1146/annurev.physiol.64.081501.160008.

Peng, Junmin, Myung Jong Kim, Dongmei Cheng, Duc M. Duong, Steven P. Gygi, and Morgan Sheng. 2004. “Semiquantitative Proteomic Analysis of Rat Forebrain Postsynaptic Density Fractions by Mass Spectrometry.” Journal of Biological Chemistry 279 (20): 21003–11. https://doi.org/10.1074/jbc.M400103200.

Rizzoli, Silvio O., and William J. Betz. 2005. “Synaptic Vesicle Pools.” Nature Reviews Neuroscience 6 (1): 57–69. https://doi.org/10.1038/nrn1583.

Roy, Marcia, Oksana Sorokina, Colin McLean, Silvia Tapia-González, Javier DeFelipe, J. Armstrong, and Seth Grant. 2018. “Regional Diversity in the Postsynaptic Proteome of the Mouse Brain.” Proteomes 6 (3): 31. https://doi.org/10.3390/proteomes6030031.

Sheng, M., and E. Kim. 2011. “The Postsynaptic Organization of Synapses.” Cold Spring Harbor Perspectives in Biology 3 (12): a005678–a005678. https://doi.org/10.1101/cshperspect.a005678.

Sheng, Morgan, and Casper C. Hoogenraad. 2007. “The Postsynaptic Architecture of Excitatory Synapses: A More Quantitative View.” Annual Review of Biochemistry 76 (1): 823–47. https://doi.org/10.1146/annurev.biochem.76.060805.160029.

Torchiano, Marco. 2016. Effsize - A Package For Efficient Effect Size Computation. Zenodo. https://doi.org/10.5281/ZENODO.1480624.

Turrigiano, G. 2012. “Homeostatic Synaptic Plasticity: Local and Global Mechanisms for Stabilizing Neuronal Function.” Cold Spring Harbor Perspectives in Biology 4 (1): a005736–a005736. https://doi.org/10.1101/cshperspect.a005736.

Turrigiano, Gina G. 2008. “The Self-Tuning Neuron: Synaptic Scaling of Excitatory Synapses.” Cell 135 (3): 422–35. https://doi.org/10.1016/j.cell.2008.10.008.

Turrigiano, Gina G., Kenneth R. Leslie, Niraj S. Desai, Lana C. Rutherford, and Sacha B. Nelson. 1998. “Activity-Dependent Scaling of Quantal Amplitude in Neocortical Neurons.” Nature 391 (6670): 892–96. https://doi.org/10.1038/36103.

Turrigiano, Gina G, and Sacha B Nelson. 2000. “Hebb and Homeostasis in Neuronal Plasticity.” Current Opinion in Neurobiology 10 (3): 358–64. https://doi.org/10.1016/S0959-4388(00)00091-X.

Turrigiano, Gina G., and Sacha B. Nelson. 2004. “Homeostatic Plasticity in the Developing Nervous System.” Nature Reviews Neuroscience 5 (2): 97–107. https://doi.org/10.1038/nrn1327.

Villa, Katherine L., Kalen P. Berry, Jaichandar Subramanian, Jae Won Cha, Won Chan Oh, Hyung-Bae Kwon, Yoshiyuki Kubota, Peter T.C. So, and Elly Nedivi. 2016. “Inhibitory Synapses Are Repeatedly Assembled and Removed at Persistent Sites In Vivo.” Neuron 89 (4): 756–69. https://doi.org/10.1016/j.neuron.2016.01.010.

Warnes, Gregory R., Ben Bolker, Lodewijk Bonebakker, Robert Gentleman, Wolfgang Huber, Andy Liaw, Thomas Lumley, et al. 2020. Gplots: Various R Programming Tools for Plotting Data. https://CRAN.R-project.org/package=gplots.

Wickham, Hadley. 2016. Ggplot2: Elegant Graphics for Data Analysis. Springer-Verlag New York. https://ggplot2.tidyverse.org.

Wilke, Claus O. 2018. Ggridges: Ridgeline Plots in “Ggplot2.” https://CRAN.R-project.org/package=ggridges.

Wing, Max Kuhn Contributions from Jed, Steve Weston, Andre Williams, Chris Keefer, Allan Engelhardt, Tony Cooper, Zachary Mayer, et al. 2019. Caret: Classification and Regression Training. https://CRAN.R-project.org/package=caret.

Zavalin, Andre, Junhai Yang, Kevin Hayden, Marvin Vestal, and Richard M. Caprioli. 2015. “Tissue Protein Imaging at 1 Mm Laser Spot Diameter for High Spatial Resolution and High Imaging Speed Using Transmission Geometry MALDI TOF MS.” Analytical and Bioanalytical Chemistry 407 (8): 2337–42. https://doi.org/10.1007/s00216-015-8532-6.

Zhou, Mu, Feixue Liang, Xiaorui R Xiong, Lu Li, Haifu Li, Zhongju Xiao, Huizhong W Tao, and Li I Zhang. 2014. “Scaling down of Balanced Excitation and Inhibition by Active Behavioral States in Auditory Cortex.” Nature Neuroscience 17 (6): 841–50. https://doi.org/10.1038/nn.3701.

Zhu, Fei, Mélissa Cizeron, Zhen Qiu, Ruth Benavides-Piccione, Maksym Kopanitsa, Nathan G. Skene, Babis Koniaris, et al. 2018. “Architecture of the Mouse Brain Synaptome.” Neuron 99 (4): 781–799.e10. https://doi.org/10.1016/j.neuron.2018.07.007.

